# Gata3 is detrimental to regulatory T cell function in autoimmune diabetes

**DOI:** 10.1101/2023.03.18.533297

**Authors:** Badr Kiaf, Kevin Bode, Cornelia Schuster, Stephan Kissler

## Abstract

Regulatory T cells (Tregs) protect against autoimmunity. In type 1 diabetes (T1D), Tregs slow the progression of beta cell autoimmunity within pancreatic islets. Increasing the potency or frequency of Tregs can prevent diabetes, as evidenced by studies in the nonobese diabetic (NOD) mouse model for T1D. We report herein that a significant proportion of islets Tregs in NOD mice express *Gata3*. The expression of *Gata3* was correlated with the presence of IL-33, a cytokine known to induce and expand Gata3^+^ Tregs. Despite significantly increasing the frequency of Tregs in the pancreas, exogenous IL-33 was not protective. Based on these data, we hypothesized that Gata3 is deleterious to Treg function in autoimmune diabetes. To test this notion, we generated NOD mice with a Treg-specific deletion of *Gata3*. We found that deleting *Gata3* in Tregs strongly protected against diabetes. Disease protection was associated with a shift of islet Tregs toward a suppressive CXCR3^+^Foxp3^+^ population. Our results suggest that islet Gata3^+^ Tregs are maladaptive and that this Treg subpopulation compromises the regulation of islet autoimmunity, contributing to diabetes onset.

## Introduction

The autoimmune attack on pancreatic beta cells that underlies type 1 diabetes (T1D) is driven by T cells (Clark et al., 2017). A majority of the genes associated with T1D by genome-wide association studies relate to T cell function (Wicker et al., 2005). In particular, a number of diabetes risk gene variants affect the function of Foxp3^+^ regulatory T cells (Tregs)(Yamanouchi et al., 2010; Gerold et al., 2011; Nowakowska and Kissler, 2016). A role for Tregs in autoimmune diabetes is exemplified in their ability to prevent or halt disease in animal models. For instance, the administration of low-dose interleukin-2 (IL-2) to NOD mice, the pre-eminent mouse model for T1D, causes Treg expansion (Grinberg-Bleyer et al., 2010). This expansion can prevent disease in pre-diabetic mice and even revert hyperglycemia in newly diabetic mice (Grinberg-Bleyer et al., 2010). In turn, the depletion of Treg cells accelerates diabetes onset in experimental models for T1D (Feuerer et al., 2009; Watts et al., 2021). Collectively, these observations show that Tregs play a central role in autoimmune diabetes. Based on their ability to modulate disease, Tregs have become a target for therapeutic intervention. Low-dose IL-2, that was effective in treating autoimmune diabetes in mice, has been tested in clinical trials to increase Tregs in patients with T1D (Todd et al., 2016; Hartemann et al., 2013).

Conventional CD4^+^ T cells do not express Foxp3 and can instead be classified by their expression of the transcription factors T-bet, Gata3 and RORγt. These transcription factors were initially thought to confer effector functions only to Th1, Th2 and Th17 cells, respectively. However, more recent studies demonstrated that Tregs can express the same transcription factors whilst retaining suppressive capacity (Levine et al., 2017; Yu et al., 2015). It is now believed that Treg subpopulations adapt to their local environment and acquire more specialized functions by also expressing T-bet, Gata3 or RORγt. The expression of these transcription factors is largely mutually exclusive (Wohlfert et al., 2011; Schiering et al., 2014; Zhu et al., 2004; Hwang et al., 2005). This may be particularly important to induce the expression of Th-like chemokine receptors that allow Tregs to co-localize with their effector T cell counterparts (Campbell, 2015). This notion has implications for the phenotype of Tregs in inflamed tissues, where effective regulation requires Tregs to adapt to the type of inflammatory cells that are present.

Treg heterogeneity has been studied in most depth at mucosal surfaces. RORγt^+^ Tregs were shown to be abundant in the colon and to control gut immunity (Sefik et al., 2015; Ohnmacht et al., 2015). Gata3^+^ Tregs were also found to be enriched at barrier sites including the skin, lungs and intestine (Wohlfert et al., 2011; Schiering et al., 2014; Harrison et al., 2019). Various studies have now demonstrated that these specialized Treg populations fill a functional niche to regulate mucosal immunity. The absence of the Gata3^+^ subset was shown to exacerbate colitis (Wohlfert et al., 2011; Schiering et al., 2014) and to cause skin inflammation (Harrison et al., 2019). Gata3 has also been associated with so-called tissue Tregs found in adipose tissue and kidney where the cells contributes to tissue repair (Muñoz-Rojas and Mathis, 2021; Sakai et al., 2021). In contrast, the composition of Tregs in the pancreas during autoimmune diabetes has not been studied as extensively, with the notable exception of one publication that reported on the strong suppressive role of Cxcr3^+^ T-bet^+^ Tregs in NOD mice (Tan et al., 2016).

We previously described that peripherally-induced Tregs (pTregs) contribute to immune regulation in autoimmune diabetes (Schuster et al., 2018). pTregs can be distinguished from thymus-derived Tregs by several differentially expressed markers including the transcription factor Helios (Thornton et al., 2010). In the course of our studies, we measured the expression of lineage-associated transcription factors in pancreatic Tregs. We made the surprising discovery that a majority of Tregs in the islets of NOD mice express Gata3. This novel observation prompted us to ask what role Gata3^+^ Tregs may play in islet autoimmunity.

## Results and Discussion

### Gata3^+^Foxp3^+^Tregs are a previously unappreciated feature of islet autoimmunity

Prompted by our earlier observation that pTregs play a role in the NOD model for type 1 diabetes (Schuster et al., 2018), we sought to further characterize the composition of the Treg population within pancreatic islets. Unexpectedly, we observed that many islet-infiltrating Tregs expressed the transcription factor Gata3, whilst T-bet^+^ and RORγt^+^ Tregs were infrequent (Fig. 1A). A role for Gata3^+^ Tregs had previously been described in the colon (Schiering et al., 2014) and visceral fat (Cipolletta et al., 2012), but this subset had not yet been associated with islet autoimmunity.

**Figure 1.**
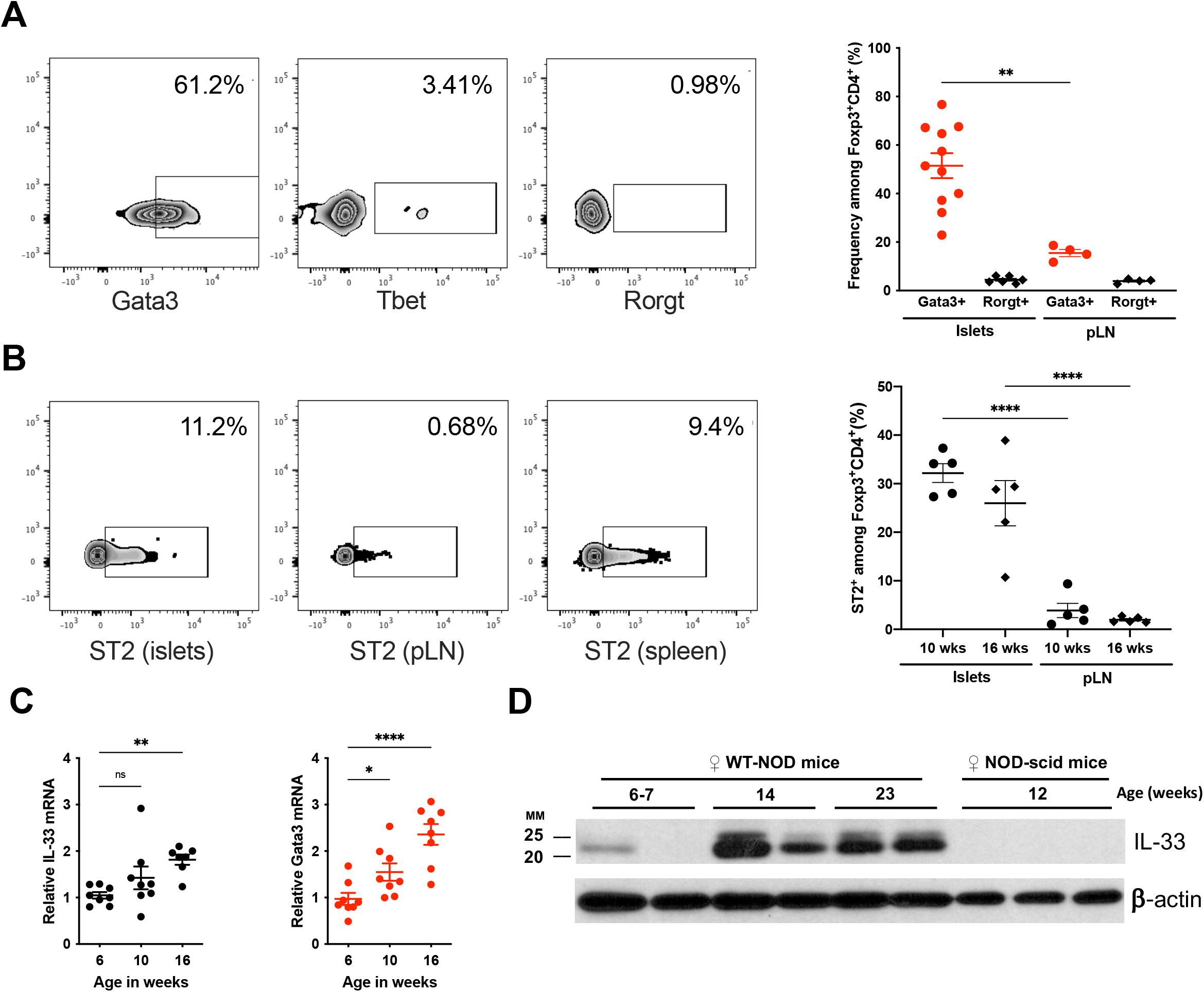
Gata3^+^Foxp3^+^Tregs are a previously unappreciated feature of islet autoimmunity. **A:** Representative flow cytometry for Gata3, T-bet and Rorγt expression gated on Foxp3^+^CD4^+^ T cells (left), and frequency of Gata3^+^ cells among Foxp3^+^CD4^+^ T cells in islets and pancreatic lymph nodes (pLN) of 10-12 week-old female NOD mice (right). The frequency of Rorγt^+^ cells is shown for comparison. (n=4-11/group) **B:** Representative flow cytometry for ST2 expression on the surface of Foxp3^+^CD4^+^ T cells in islets, pLN and spleen (left). Summary data are shown for ST2 expression in islet and pLN Tregs in 10- and 16-week-old female NOD mice (n=5/group). **C:** Quantitative PCR measurement for *Il-33* and *Gata3* mRNA in islets from female NOD mice at the indicated age (n=7-8/group). **D:** Western blot measurement of IL-33 protein expression in islets from female NOD (n=2/age group) and NOD.scid (n=3) mice. ns P>0.05; *P<0.05; ** P<0.01; ****P<0.0001 (A: Mann-Whitney test, B,C: One-way ANOVA)

Powrie and colleagues reported that the alarmin IL-33 can induce and expand Gata3^+^ Tregs (Schiering et al., 2014). IL-33 signaling requires cell surface expression of ST2 (Il1rl1) that combines with the IL1R-accessory protein (IL1-RaP) to form the IL-33 receptor. We found that ST2 expression was more frequent in islet Tregs than in the nearby pancreatic lymph node (PLN) (Fig. 1B), suggesting that islet Tregs are equipped to respond to IL-33.

A study of pancreatic islets in a high-fat diet model for type 2 diabetes had shown that IL-33 can be released within stressed islets (Dalmas et al., 2017). We hypothesized that immune infiltration of pancreatic islets during autoimmune diabetes may similarly cause IL-33 release which could in turn induce or expand Gata3^+^ Tregs. Consistent with this hypothesis, the levels of both *Il33* and *Gata3* mRNA increased in islets as NOD mice progressed towards diabetes onset (Fig. 1C). We confirmed that IL-33 protein was abundant in islets at the late pre-diabetic stage, but not in younger NOD mice (Fig. 1D). Notably, IL-33 was undetectable in islets from immuno-deficient NOD.SCID mice that are devoid of islet inflammation and do not develop diabetes (Fig. 1D). Together, these data suggest that IL-33 released upon immune infiltration of pancreatic islets may shape the Treg compartment and increase the frequency of Gata3^+^Foxp3^+^ T cells.

### IL-33 increases the frequency of Gata3^+^Foxp3^+^ Tregs within islets

To directly test if IL-33 can induce or expand Gata3^+^ Tregs in the pancreas, we administered recombinant IL-33 to pre-diabetic NOD mice over a 7-day period. Exogenous IL-33 caused a significant increase in the frequency of Tregs in the spleen as described previously (Schiering et al., 2014), but also in the pancreas (Fig. 2A). Notably, IL-33 increased the frequency of Tregs that expressed ST2 and Gata3. The elevated Treg frequency persisted for at least 3 weeks after treatment with IL-33 (Fig. 2B).

**Figure 2.**
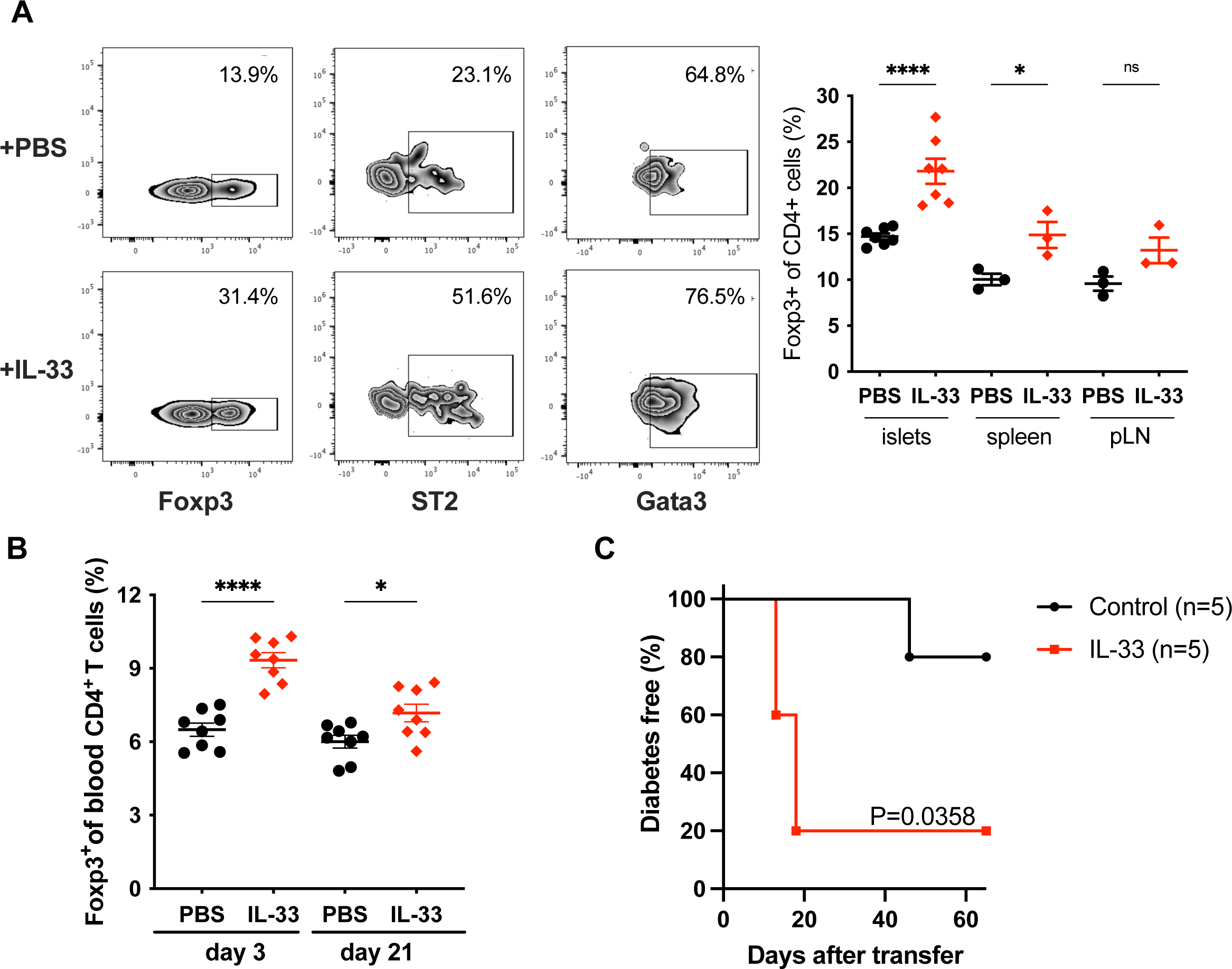
IL-33 increases the frequency of Gata3^+^Foxp3^+^ Tregs within islets but is not protective. **A:** Female NOD mice were injected with 1μg rIL-33/day or vehicle (PBS) for 7 days. Left: Representative flow cytometry data measuring Foxp3 expression (gated on CD4^+^ T cells) and Gata3 or ST2 expression (gated on Foxp3^+^CD4^+^ T cells). Right: Frequency of Foxp3^+^CD4^+^ T cells in the islets, pLN and spleen of mice treated or not with rIL-33 (n=3-7/group). **B:** Frequency of Foxp3^+^CD4^+^ T cells in the blood of mice treated with or without rIL-33 at the indicated time after the first injection. **C:** Diabetes frequency in NOD.scid mice injected with splenocytes from spontaneously diabetic NOD mice and injected or not with rIL-33 (1 μg/day for 7 days), n=5/group. ns P>0.05; *P<0.05; ****P<0.0001 (A,B: One-way ANOVA). Exact P value for Log-rank (Mantel-Cox) test is shown in C.

Expansion of Tregs by low-dose IL-2 is protective in NOD mice (Grinberg-Bleyer et al., 2010). To test if expansion of Tregs by IL-33 would have a similar protective effect, we induced diabetes by adoptive transfer with or without concomitant recombinant IL-33 treatment. We found that IL-33 administration did not protect against diabetes in this setting (Fig. 2C), despite the fact that this treatment regimen vastly increased the frequency of pancreatic Tregs.

### Treg-specific deletion of Gata3 protects against autoimmune diabetes

Based on our observation that Gata3^+^ Tregs were frequent in islets of late pre-diabetic mice and that the induction of this subset with exogenous IL-33 was not protective, we hypothesized that Gata3 expression may be a maladaptive response of Tregs in the inflamed islet microenvironment. To test this hypothesis, we generated NOD mice that harbor Tregs unable to express *Gata3*. We backcrossed a floxed *Gata3* allele (Amsen et al., 2007) from the C57BL/6 strain into the NOD background for 10 generations. We then bred NOD *Gata3*^fl/+^ mice with NOD mice that express the *cre* recombinase under the control of the *Foxp3* promoter (NOD *Foxp3-Cre*, (Zhou et al., 2008)). Resulting off-spring were intercrossed to generate cohorts of *Foxp3-cre*^+^ *Gata3*^fl/fl^ and *Foxp3-cre*^+^ *Gata3*^fl/+^ NOD mice (Fig. 3A). Remarkably, the Treg-specific deletion of *Gata3* afforded very strong protection against diabetes (Fig. 3B). Only 12% of the *Foxp3-cre*^+^ *Gata3*^fl/fl^ animals developed diabetes, while 50% of the control *Foxp3-cre*^+^ *Gata3*^fl/+^ mice did (P=0.0048, Log-rank test).

**Figure 3.**
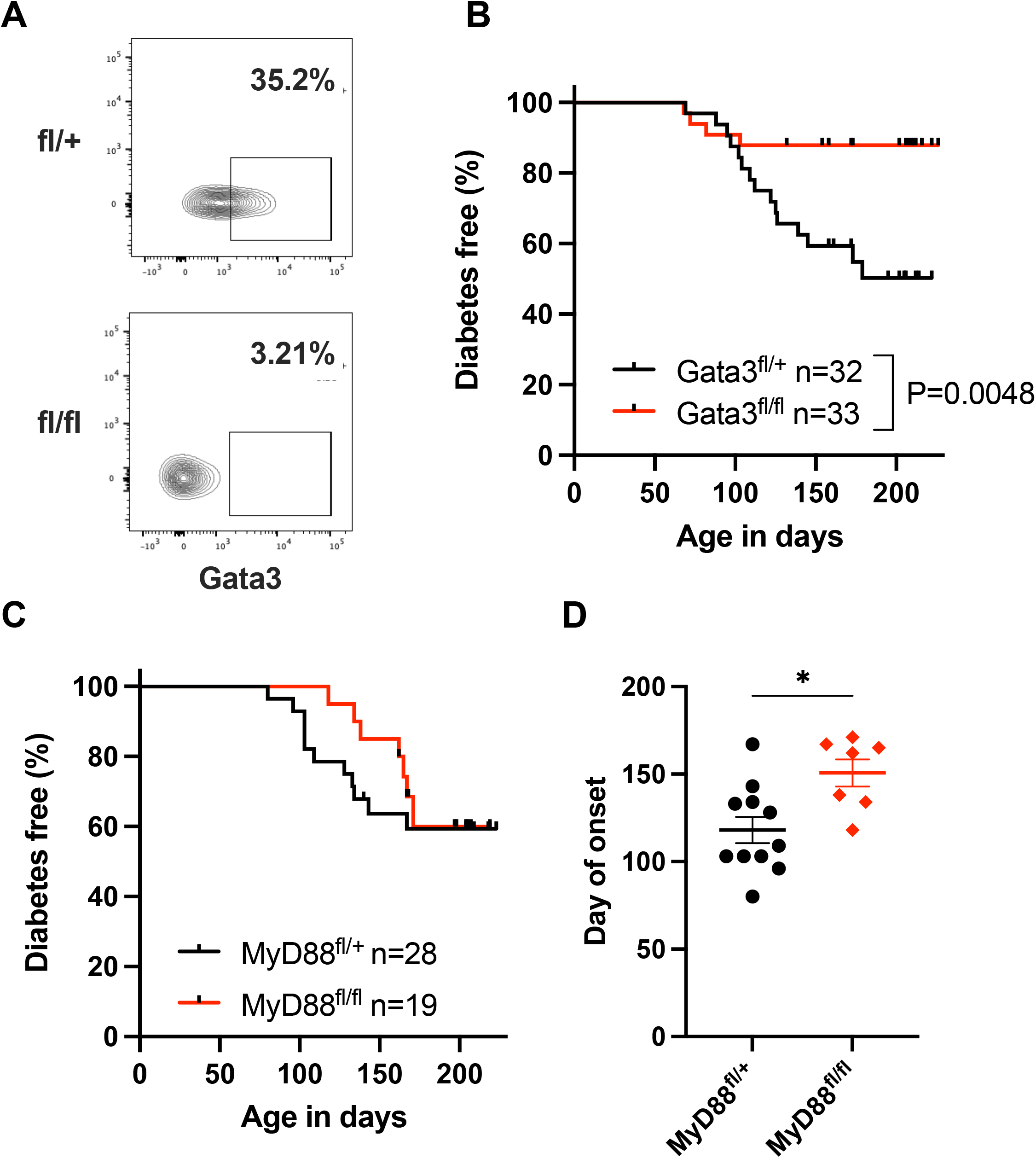
Treg-specific deletion of Gata3 protects against autoimmune diabetes. **A:** Representative flow cytometry data for Gata3 expression in pancreatic Tregs from Gata3^fl/+^Foxp3^Cre^ (top) and Gata3^fl/fl^Foxp3^Cre^ mice and showing effective *Gata3* deletion in Tregs. **B:** Diabetes frequency in cohorts of female Gata3^fl/+^ Foxp3^Cre^ (n=32) and Gata3^fl/fl^Foxp3^Cre^ (n=33) NOD mice followed over a period >200 days. P=0.0048, Log-rank (Mantel-Cox) test. **C**,**D:** Diabetes frequency (C) and day of disease onset (D) in cohorts of female MyD88^fl/+^Foxp3^Cre^ (n=28) and MyD88^fl/fl^Foxp3^Cre^ (n=19) NOD mice followed over a period >200 days. *P<0.05 (Mann-Whitney test).

We next sought to test if IL-33 signaling in Tregs had a similar critical role in diabetes pathogenesis. The ST2/IL1-RaP complex signals via the MyD88 adapter. To abrogate this signaling pathway, we introduced a floxed allele for *MyD88* (Hou et al., 2008) into NOD mice by again backcrossing for 10 generations. NOD *MyD88*^fl/+^ mice were then bred with NOD *Foxp3-cre* animals to generate cohorts of *Foxp3-cre*^+^ *MyD88*^fl/fl^ and *Foxp3-cre*^+^ *MyD88*^fl/+^ mice. The deletion of *MyD88* in the Treg compartment did not change the overall frequency of diabetes (Fig. 3C). However, *MyD88* deficiency in Tregs delayed disease onset by more than 50 days (median onset: *MyD88*^fl/+^ 109 days; *MyD88*^fl/fl^ 162 days, P=0.014, two-tailed Mann-Whitney test) (Fig. 3D). The data suggest that IL-33 plays a partial role in shaping islet T cell regulation, but that loss of IL-33 signaling in Tregs does not fully recapitulate the strong protective phenotype of *Gata3* deficiency.

### Treg-specific deletion of Gata3 increases the frequency of highly suppressive Cxcr3^+^ Tregs

To understand how the deletion of *Gata3* impacted the composition of islet Tregs, we isolated islets from the pancreas of 12-week old pre-diabetic *Foxp3-cre*^+^*Gata3*^fl/fl^ and control NOD mice. We sorted infiltrating T cells by flow cytometry and analyzed purified T cells by single-cell RNA sequencing (scRNAseq). UMAP analysis identified 20 distinct cell clusters (Fig. 4A), of which four had a Treg-like transcriptional profile (clusters 4, 8, 13 and 14). Cells in cluster 14 did not express Foxp3 (Fig. 4B) and instead had elevated levels of transcripts characteristics for Tr1 regulatory CD4^+^ T cells, including *Lag3, Maf, Il10* and *Il21* (Pot et al., 2009) (data not shown). We focused our analysis on clusters 4, 8 and 13, all of which expressed *Foxp3* and *Ikzf2* that encodes Helios (Fig. 4B). Notably, cells in cluster 4 were characterized by elevated levels of *Ikzf2, Neb, Areg, Ccr5, Tnfrsf9* and *Ccl5*, which have been associated with tissue-Tregs (Muñoz-Rojas and Mathis, 2021)(Fig. 4C). In contrast, cells in cluster 13 expressed high levels of *Cxcr3, Tcf7* and *Lef1* that mark Cxcr3^+^Tbet^+^ Tregs that are highly suppressive (Xing et al., 2019)(Fig. 4C) and were shown to be critical in the control of autoimmune diabetes (Tan et al., 2016).

**Figure 4.**
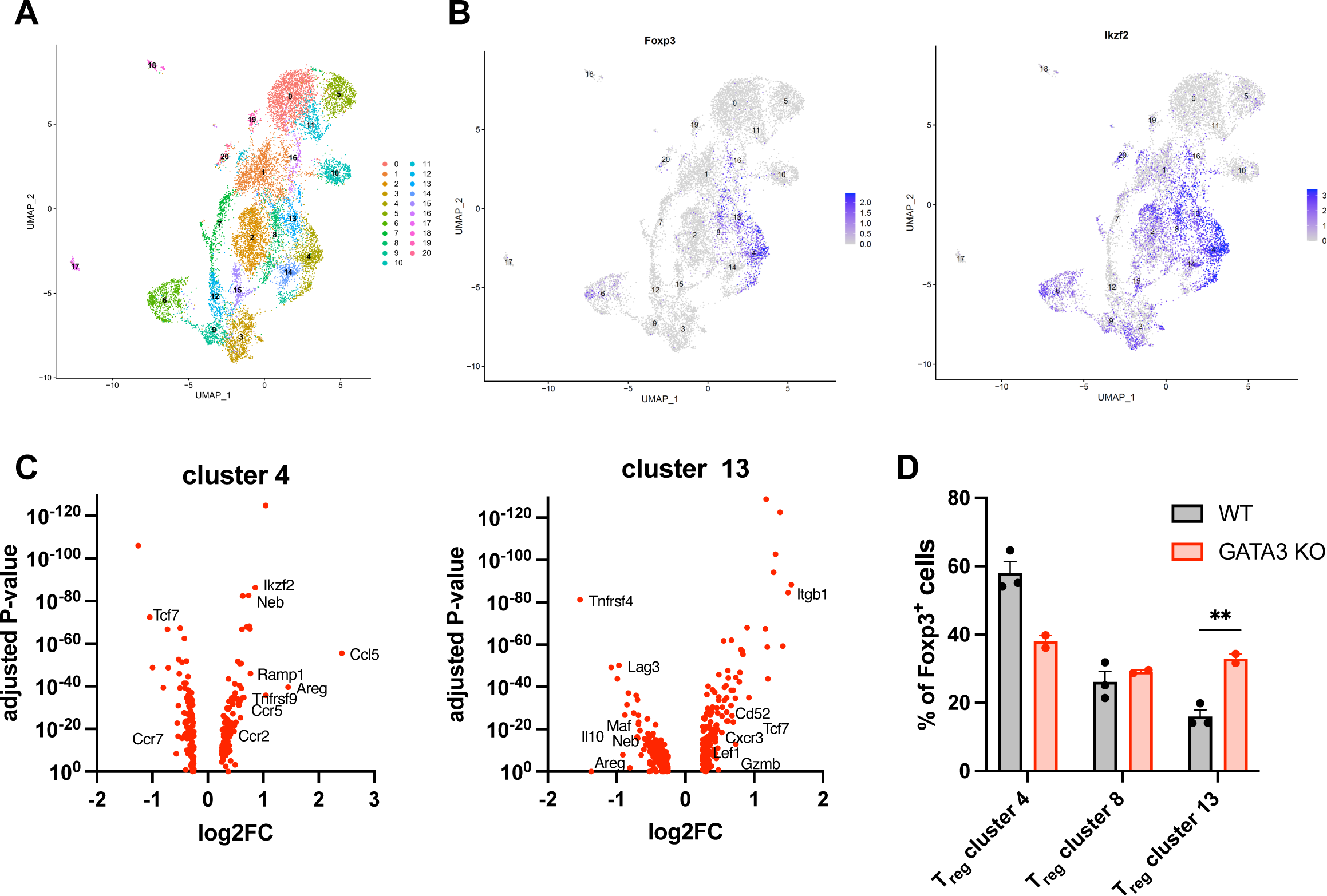
Treg-specific deletion of Gata3 modifies the composition of islets Tregs. **A:** UMAP representation of islet-infiltrating T cells sorted by flow cytometry (CD45^+^TCRβ^+^PI^-^) from pre-diabetic 12-week old NOD mice, identifying 20 clusters. **B:** Expression of *Foxp3* (left panel) and *Ikzf2* (right panel) across all cell clusters. **C:** Volcano plots for genes up- or down-regulated within clusters 4 (left panel) and 13 (right panel) in comparison to all Foxp3^+^ cells. The most relevant genes associated with previously described Treg subpopulations are denoted next to the corresponding data points. **D:** Frequency of cells within clusters 4, 8 and 13 relative to the total Foxp3^+^CD4^+^ T cells population identified in these three Treg clusters. Data represent cells isolated from 2 *Foxp3-Cre*^+^*Gata3*^fl/fl^ and 3 WT mice. ^**^P=0.009 (two-sided unpaired t-test).

Due to the technical limitations of scRNAseq that does not capture the entirety of the transcriptome, the expression of *Gata3* and *Tbet* in these populations could not be ascertained. Notwithstanding, the data are strongly suggestive of cluster 4 identifying a tissue-Treg-like population and of cluster 13 representing the highly suppressive Cxcr3^+^Tbet^+^ Tregs described in the islet of NOD mice previously (Tan et al., 2016). Importantly, the Treg-specific deletion of *Gata3* significantly increased the frequency of suppressive Tregs in cluster 13 while diminishing the frequency of tissue-Treg cells in cluster 4 (Fig. 4D). We propose that *Gata3* deletion skews the fate of islet Tregs away from a reparative tissue-Treg phenotype in favor of a more suppressive transcriptional program. Preventing islet Tregs from acquiring a maladaptive Gata3-dependent phenotype may improve regulation of islet autoimmunity and protect against diabetes by increasing the frequency of Tregs that are better equipped to suppress autoreactive effector T cells.

## Conclusions

In this study, we investigated the heterogeneity of Tregs in the inflamed islet environment of NOD mice that develop autoimmune diabetes. We discovered that islet Tregs comprise a large fraction of Gata3^+^ cells that express ST2, a component of the IL-33R, and are responsive to the alarmin IL-33. Counterintuitively, increasing the frequency of Gata3^+^ Tregs by IL-33 administration was not protective. We speculated that Gata3 expression in islet Tregs may be maladaptive and that Gata3^+^ Tregs may be ill-equipped to control islet autoimmunity. In support of this hypothesis, Treg-conditional deletion of Gata3 was highly protective against the development of autoimmune diabetes. Abrogating IL-33 signaling in Tregs by conditional deletion of MyD88 also delayed the onset of disease, though it did not fully replicate the protection afforded by Treg-specific Gata3 deletion. We propose that the expression of Gata3 in islet Tregs, driven in part by local IL-33 release, hinders the regulation of autoimmunity in T1D and contributes to disease progression.

The expression of Gata3 in Foxp3^+^ Tregs has been associated with a tissue-Treg phenotype (Muñoz-Rojas and Mathis, 2021). Reports describing tissue Tregs in various organs have provided evidence for a divergent role of these cells relative to the classical immune suppressive function associated with Foxp3^+^CD4^+^ T cells. In the context of tissue damage, Tregs can adopt a reparative function (Arpaia et al., 2015; Burzyn et al., 2013). We propose that islet Tregs involved in the autoimmune response in T1D are diverted towards a tissue-repair phenotype by local factors that include IL-33. This diversion towards a reparative phenotype would weaken immune regulation in the inflamed islet and accelerate disease onset. This could explain why autoimmune diabetes develops in NOD mice despite a high frequency of Tregs within the pancreas (Mellanby et al., 2007; Kaur et al., 2010). Disease progresses not because of a numerical deficiency of islet Tregs but rather because of their functional maladaptation. Consistent with this notion, the Treg-specific deletion of T-bet was reported to have the opposite effect and to markedly increase the risk of diabetes (Tan et al., 2016). In this setting, the ablation of T-bet compromised the functionality of islet Tregs, suggesting that T-bet^+^ Foxp3^+^ T cells, but not Gata3^+^ Foxp3^+^ cells, are effective regulators of islet autoimmunity. The balance between an immune suppressive versus a tissue reparative Treg phenotype may determine whether or not autoreactive T cells are kept in check in the inflamed pancreas.

Of interest, a recent mass cytometry analysis identified an increase in GATA3^+^ Tregs in the blood of children at high risk of T1D (Barcenilla et al., 2019). Of all the immune populations examined in this study, only GATA3^+^FOXP3^+^CD4^+^ T cells and natural killer (NK) cells differed in their frequencies between individuals at high risk and the control group. While little is known of the function of human GATA3^+^ Tregs, this observation is intriguing and suggests a possible role for Treg maladaptation in human T1D.

In sum, we propose that Treg maladaptation within the islet environment contributes to the pathogenesis of autoimmune diabetes. Diverting Tregs toward a T-bet-dependent immune suppressive phenotype and away from a Gata3-driven tissue repair program may hold therapeutic potential, particularly for future interventions utilizing CAR-Tregs (Yeh et al., 2017; Tenspolde et al., 2019; Radichev et al., 2020).

## Acknowledgements

This research was supported by the Harvard Stem Cell Institute with funds from a collaborative program with the Qatar Biomedical Research Institute / Hamad Bin Khalifa University. The research was also supported by Joslin core facilities funded in part by a Diabetes Research Center award from NIDDK (P30DK036836). B.K., K. B. and C.S. were recipients of postdoctoral fellowships from the Mary K. Iacocca Foundation.

## Author contributions

C.S. performed experiments and made the initial observation underlying this study. B.K. developed the concept of Treg maladaptation, designed and performed experiments, interpreted the data and helped in the preparation of the manuscript. K.B. performed scRNAseq experiments, analyzed the data and helped in the preparation of the manuscript. S.K. conceived the study, designed experiments, interpreted data and wrote the manuscript. All authors edited the manuscript and approved the final version.

## Methods

### Mice and diabetes measurements

GATA3^fl/fl^ (Jackson lab, #028103) and MyD88^fl/fl^ (Jackson lab, #009108) NOD mice were generated by backcrosses (>12 times) GATA3^fl/fl^ C57BL/6J and MyD88^fl/fl^ C57BL/6J on the NOD background.

Breeding was performed between GATA3^fl/fl^ or GATA3^fl/wt^ and MyD88^fl/fl^ or MyD88^fl/wt^ NOD mice with FoxP3^Cre/wt^ or FoxP3^Cre/wt^ NOD mice (Jackson lab, #008694) to generate mice with GATA3 or MyD88 Treg specific knock-out and littermate controls. All mice were bred under specific pathogen-free conditions. Disease studies were performed with age-matched, contemporary cohorts of mice. Onset of diabetes was monitored by weekly measurements of glycemia using ACCU-CHEK, Roche. Mice with two consecutive readings > 250 mg/dL were considered diabetic.

To study the effect of IL-33 on Treg cells, NOD female mice were injected intraperitoneally (i.p.) with recombinant mouse IL-33 (0.5-1ug/mice/day, in 100uL of PBS; BioLegend: 580508) or vehicle only (PBS) daily for 7 days.

To induce diabetes, 10^7^ naïve CD4+ T cells isolated from the spleen of Th1.1 congenic BDC2.5 NOD mice were injected intravenously (i.v.) into sex matched Th1.2 congenic NOD^scid^ mice. Mice were then injected i.p. with rmIL-33 (0.5-1ug/mice/day, in 100uL of PBS; BioLegend: 580508) for 7 days or vehicle (PBS only). Mice were tested weekly and considered diabetic after two consecutive blood glycemia readings of > 250 mg/dL using an ACCU-CHEK blood glucose monitor (Roche).

### Lymphocytes isolation

Blood was collected from the tail vein in capillary EDTA-coated tubes and lymphocytes were stained for flow cytometry after red blood cell lysis. Single cell suspensions were prepared from spleen and lymph nodes by mechanical disruption of tissue followed by red blood cell lysis using ACK buffer.

Islet isolations were performed using intraductal collagenase digestion. Briefly, while mice were under terminal anesthesia, the common bile duct was clamped and the pancreatic duct was perfused with 3-5 mL solution of collagenase P in HBSS-1% HEPES (Roche). The pancreas was then harvested and transferred to a 50mL conical tube containing 5mL of collagenase P solution and kept on ice until all organs were collected. Pancreases were digested at 37°C for 20 min. The digestion was stopped by adding HBSS-10% FBS, followed by 2 washes. Islets were filtered through a strainer, handpicked and counted. Alternatively, the islets were separated by a density gradient (Histopaque-1077; Sigma-Aldrich, 10mL per pancreas, 2000rpm, 20 minutes, without brakes). For immune cells isolation, islets were incubated (37°C for 6 min) in a non-enzymatic cell dissociation solution (Sigma-Aldrich; #C5914-100ML) then vortexed to disrupt the islets.

### Quantitative RT-PCR

RNA was extracted from tissues or purified cells using the the NucleoSpin® RNA Plus Kit (Macherey-Nagel). Complementary DNAs were synthesized using the SuperScript™ III First-Strand Synthesis System Kit (Invitrogen) or the AzuraQuant™ cDNA Synthesis Kit (Azura Genomics). Quantitative PCR analysis was performed using the Power SYBR™ Green PCR Master Mix (Applied Biosystems) or the AzuraQuant™ Green Fast qPCR Mix HiRox (Azura Genomics). Beta actin and GAPDH were used for normalization to quantify relative mRNA expression levels. Relative changes in mRNA expression were calculated using the comparative cycle methods 2−ΔCt or 2−ΔΔCt. The Primers are as following (GATA-3, Fw: GTCATCCCTGAGCCACATCT, Rv: AGGGCTCTGCCTCTCTAACC; IL-33, Fw: ATGGGAAGAAGCTGATGGTG, Rv: CCGAGGACTTTTTGTGAAGG)

### Flow Cytometry

Flow cytometry measurements were performed using a LSRII or LSR Fortessa instrument (BD Biosciences). Data were analyzed with FlowJo software (BD Biosciences). Fluorescently conjugated antibodies were purchased from Biolegend, eBioscience/Thermo Fisher Scientific and BD Biosciences. Intracellular staining was performed using a Cytofix/Cytoperm Plus Kit (BD Biosciences) or a FoxP3/Transcription Factor Staining Buffer Set (eBioscience/Thermo Fisher Scientific). Surface staining was performed in PBS +2% FBS. For exclusion of dead cells, Zombie Fixable Viability Dyes were used (Biolegend).

### Western blotting

Isolated islets from female NOD mice of NOD Scid mice were digested in 100uL of digestion buffer (1X Cell Lysis Buffer (Cell Signaling Technology)) supplemented with protease inhibitors (cOmplete™, Mini, EDTA-free Protease Inhibitor Cocktail, Roche) and 1mM PMSF (Cell Signaling Technology).

10 ug of the protein lysate were mixed with 4X Laemmli buffer (Bio-Rad) supplemented with 2-mercaptoethanol (Sigma-Aldrich) at a 3:1 (sample:buffer) ratio were denatured at 95°C for 5 minutes before loading onto a 15% SDS-PAGE gel, followed by a transfer onto a nitrocellulose membrane (Bio-Rad). Proteins were detected using Rat anti-IL-33 (R&D Systems) and Rabbit β-Actin (Cell Signaling Technology) antibodies followed by HRP-conjugated secondary antibodies (anti-Rabbit or anti-Rat antibodies (Cell Signaling Technology).

### Single cell RNA sequencing (scRNAseq) of islet-infiltrating T cells

Islets were isolated and purified as described above from 12-week old non-diabetic female WT or Foxp3^Cre^GATA3^fl/fl^ NOD mice. Briefly, the pancreas was perfused with collagenase type V/cold Hank’s balanced salt solution (HBSS) and was immediately removed by surgical dissection. The pancreatic tissue was digested in a 37 °C water bath for 15–17 min. The digested tissue was then washed with cold HBSS three times, followed by Histopaque gradient separation. Islets were handpicked under a dissection microscope and further digested using HEPES buffered RMPI 1640 (cat # R4130-10L; Sigma-Aldrich) supplemented with 1mg/ml Collagenase D (cat # 11088858001; Sigma-Aldrich), 20 μg/ml DNase I (cat # EN0521; Thermo Fisher Scientific), 2% FCS and 50 μg/ml Lipopolysaccharide neutralizing agent Polymyxin B sulfate (cat # 1405-20-5; Sigma-Aldrich) for 30 min at 37°C while shaking by 750 rpm on a heating block. Single cell suspension were generated by filtering the digested islets through a 70uM strainer. Cells were collected and stained with fluorescently labeled antibodies against CD45 PE-Cy 7 (clone: 30-F11, cat # 103114: Biolegend), TCRβ APC (clone: H57-597, cat # 109212: Biolegend) and with the viability dye Propidium Iodide (PI, cat # R37169; Thermo Fisher Scientific). Next, islet-infiltrated T cells were FACS sorted by CD45^+^TCRβ^+^ PI^-^ (3 biological replicates for each genotype) by using a *SORP BD FACSAria*. A maximum of 10,000 cells per sample were used for scRNAseq using the Chromium Next GEM Single Cell 3’ GEM, Library & Gel Bead Kit v3.1 (cat # PN-1000213; 10 x Genomics) according to manufactures’ instructions. Illumina NovaSeq 6000 with about 1.3 billion reads total was used for sequencing the purified 3’ gene expression library.

Gene UMI counts in droplets by aligning reads to the mouse genome (mm10, GENCODE vM23/Ensembl 98) using the count function of cellranger. To distinguish between droplets containing cells RNA and ambient RNA, we used Monte Carlo simulations to compute *p*-values for the multinomial sampling transcripts from the ambient pool (Lun et al., 2019). First, we assumed that some barcodes correspond to empty droplets if their total UMI counts were at or below 50. We then called cells at a false discovery rate (FDR) of 0.1%, meaning that no more than 0.1% of our called barcodes should be empty droplets on average. The number of Monte Carlo iterations determined the lower bound for the p-values (Phipson and Smyth 2010). There were no non-significant barcodes bounded by iterations, which indicated we didn’t need to increase the number of iterations to obtain even lower p-values.

We computed the maximum contribution of the ambient solution to the expression profile for cell-containing droplets (Lun et al., 2019). Firstly, we estimated the composition of the ambient pool of RNA based on the barcodes with total UMI counts less than or equal to 50. Secondly, we computed the mean ambient contribution for each gene by scaling the ambient pool by some factor. Thirdly, we computed a p-value for each gene based on the probability of observing an ambient count equal to or below that in cell-containing droplets based on Poisson distribution. Fourthly, we combined p-values across all genes using Simes’ method (Simes, 1986). We performed this for a range of scaling factors and identify the largest factor that yields a combined p-value above threshold 0.1 so that the ambient proportions were the maximum estimations.

We performed Principal component analysis (PCA) on the transformed data. Only the top 3000 variable genes were used as input. We constructed a K Nearest Neighbor (KNN) network of the cells based on the Euclidean distance in the space defined by the top principal components (PCs) that were selected by Horn’s parallel analysis (Horn, 1965). We refined the weights of the connection between pairs of cells based on their shared overlap in their local neighborhoods *i*.*e*. Jaccard similarity (Jaccard, 1912). To cluster the cells, we applied modularity optimization techniques, i.e. the Louvain algorithm (Blondel et al., 2008). We removed the ambient RNA contamination from the cluster-level profiles and propagated the effect of those changes back to the individual cells using the removeAmbience function of the R package DropletUtils (Lun et al., 2019).

We identified doublets using the R package scDblFinder (Germain et al., 2022) and then removed doublets from the downstream analysis. We used the QC metrics to filter cells that had less than 1000 UMI (low-quality cells), less than 500 features (low-quality cells), and more than 10% mitochondrial UMI (dying cells). We normalized the combined data using R package sctransform (Hafemeister and Satija, 2019).

For cell cluster assignment of combined samples we constructed a K Nearest Neighbor (KNN) network of the cells based on the Euclidean distance in the space defined by the top principal components (PCs) that were selected by Horn’s parallel analysis (Horn, 1965). We refined the weights of the connection between pairs of cells based on their shared overlap in their local neighborhoods i.e. Jaccard similarity (Jaccard, 1912). To cluster the cells, we applied modularity optimization techniques, i.e. the Louvain algorithm (Blondel et al., 2008). We identified a total of 20 cell clusters. We performed UMAP analysis using the same PCs as input to the clustering analysis.

For differential expression genes (DEG) analysis between cell clusters we found markers for every cluster compared to all remaining cells using the Wilcoxon Rank Sum test, reported only ones with a log2FC threshold greater than 0.25 and expressed in more than 10% of the cells. We then made a heatmap for the top 5 up-regulated markers of each cluster (selected by log2FC). To discover the differential genes between genotypes in T_reg_ cluster, we aggregated (sum) the gene counts for each cell type in each sample and filter out low expressing genes but kept genes that have counts per million (CPM) more than 1 in at least 2 samples of a cell type. To discover the differential genes, we used edgeR, an R package for differential expression analysis of digital gene expression data (Robinson and Oshlack, 2010). We performed generalized Quasi-Likelihood F-tests to detect genes that were differentially expressed between groups in each cell type. R version 4.1.1 (2021-08-10) and the Platform: x86_64-conda-linux-gnu (64-bit) was used.

## References

Amsen, D., A. Antov, D. Jankovic, A. Sher, F. Radtke, A. Souabni, M. Busslinger, B. McCright, T. Gridley, and R.A. Flavell. 2007. Direct Regulation of Gata3 Expression Determines the T Helper Differentiation Potential of Notch. Immunity. 27:89–99. doi:10.1016/j.immuni.2007.05.021.

Arpaia, N., J.A. Green, B. Moltedo, A. Arvey, S. Hemmers, S. Yuan, P.M. Treuting, and A.Y. Rudensky. 2015. A Distinct Function of Regulatory T Cells in Tissue Protection. Cell. 162:1078–1089. doi:10.1016/J.CELL.2015.08.021.

Barcenilla, H., L. Åkerman, M. Pihl, J. Ludvigsson, and R. Casas. 2019. Mass Cytometry Identifies Distinct Subsets of Regulatory T Cells and Natural Killer Cells Associated With High Risk for Type 1 Diabetes. Front. Immunol. 10. doi:10.3389/fimmu.2019.00982.

Blondel, V., J.-L. Guillaume, R. Lambiotte, and E. Lefebvre. 2008. Fast unfolding of communities in large networks. J. Stat. Mech. P10008.

Burzyn, D., W. Kuswanto, D. Kolodin, J.L. Shadrach, M. Cerletti, Y. Jang, E. Sefik, T.G. Tan, A.J. Wagers, C. Benoist, and D. Mathis. 2013. A special population of regulatory T cells potentiates muscle repair. Cell. 155. doi:10.1016/J.CELL.2013.10.054.

Campbell, D.J. 2015. Control of Regulatory T Cell Migration, Function, and Homeostasis. J. Immunol. 195:2507–2513. doi:10.4049/jimmunol.1500801.

Cipolletta, D., M. Feuerer, A. Li, N. Kamei, J. Lee, S.E. Shoelson, C. Benoist, and D. Mathis. 2012. PPAR-γ is a major driver of the accumulation and phenotype of adipose tissue Treg cells. Nature. 486:549–53. doi:10.1038/nature11132.

Clark, M., C.J. Kroger, and R.M. Tisch. 2017. Type 1 Diabetes: A Chronic Anti-Self-Inflammatory Response. Front. Immunol. 8:1898. doi:10.3389/fimmu.2017.01898.

Dalmas, E., F.M. Lehmann, E. Dror, S. Wueest, C. Thienel, M. Borsigova, M. Stawiski, E. Traunecker, F.C. Lucchini, D.H. Dapito, S.M. Kallert, B. Guigas, F. Pattou, J. Kerr-Conte, P. Maechler, J.-P. Girard, D. Konrad, C. Wolfrum, M. Böni-Schnetzler, D. Finke, and M.Y. Donath. 2017. Interleukin-33-Activated Islet-Resident Innate Lymphoid Cells Promote Insulin Secretion through Myeloid Cell Retinoic Acid Production. Immunity. 47:928-942.e7. doi:10.1016/j.immuni.2017.10.015.

Feuerer, M., Y. Shen, D.R. Littman, C. Benoist, and D. Mathis. 2009. How punctual ablation of regulatory T cells unleashes an autoimmune lesion within the pancreatic islets. Immunity. 31:654–64. doi:10.1016/j.immuni.2009.08.023.

Germain, P.L., M.D. Robinson, A. Lun, C. Garcia Meixide, and W. Macnair. 2022. Doublet identification in single-cell sequencing data using scDblFinder. F1000Research. 10. doi:10.12688/F1000RESEARCH.73600.2/DOI.

Gerold, K.D.K.D., P. Zheng, D.B.D.B. Rainbow, A. Zernecke, L.S.L.S. Wicker, and S. Kissler. 2011. The soluble CTLA-4 splice variant protects from type 1 diabetes and potentiates regulatory T-cell function. Diabetes. 60:1955–63. doi:10.2337/db11-0130.

Grinberg-Bleyer, Y., A. Baeyens, S. You, R. Elhage, G. Fourcade, S. Gregoire, N. Cagnard, W. Carpentier, Q. Tang, J. Bluestone, L. Chatenoud, D. Klatzmann, B.L. Salomon, and E. Piaggio. 2010. IL-2 reverses established type 1 diabetes in NOD mice by a local effect on pancreatic regulatory T cells. J. Exp. Med. 207:1871–8. doi:10.1084/jem.20100209.

Hafemeister, C., and R. Satija. 2019. Normalization and variance stabilization of single-cell RNA-seq data using regularized negative binomial regression. Genome Biol. 20:1–15. doi:10.1186/S13059-019-1874-1/FIGURES/6.

Harrison, O.J., J.L. Linehan, H.Y. Shih, N. Bouladoux, S.J. Han, M. Smelkinson, S.K. Sen, A.L. Byrd, M. Enamorado, C. Yao, S. Tamoutounour, F. Van Laethem, C. Hurabielle, N. Collins, A. Paun, R. Salcedo, J.J. O’Shea, and Y. Belkaid. 2019. Commensal-specific T cell plasticity promotes rapid tissue adaptation to injury. Science. 363. doi:10.1126/SCIENCE.AAT6280.

Hartemann, A., G. Bensimon, C.A. Payan, S. Jacqueminet, O. Bourron, N. Nicolas, M. Fonfrede, M. Rosenzwajg, C. Bernard, and D. Klatzmann. 2013. Low-dose interleukin 2 in patients with type 1 diabetes: a phase 1/2 randomised, double-blind, placebo-controlled trial. lancet. Diabetes Endocrinol. 1:295–305. doi:10.1016/S2213-8587(13)70113-X.

Horn, J.L. 1965. A rationale and test for the number of factors in factor analysis. Psychometrika. 30:179– 185.

Hou, B., B. Reizis, and A.L. DeFranco. 2008. Toll-like receptors activate innate and adaptive immunity by using dendritic cell-intrinsic and -extrinsic mechanisms. Immunity. 29:272–282. doi:10.1016/J.IMMUNI.2008.05.016.

Hwang, E.S., S.J. Szabo, P.L. Schwartzberg, and L.H. Glimcher. 2005. T helper cell fate specified by kinase-mediated interaction of T-bet with GATA-3. Science (80-.). 307:430–433. doi:10.1126/SCIENCE.1103336/SUPPL_FILE/HWANG_SOM.PDF.

Jaccard, P. 1912. The distribution of flora of the alpine zone. New Phytol. 11:37–50.

Kaur, S., W.L. Tan, C. Soo, C.C.H. Cheung, J. Stewart, and S. Reddy. 2010. An immunohistochemical study on the distribution and frequency of T regulatory cells in pancreatic islets of NOD mice during various stages of spontaneous and cyclophosphamide-accelerated diabetes. Pancreas. 39:1024–1033. doi:10.1097/MPA.0B013E3181DA9037.

Levine, A.G., A. Mendoza, S. Hemmers, B. Moltedo, R.E. Niec, M. Schizas, B.E. Hoyos, E. V Putintseva, A. Chaudhry, S. Dikiy, S. Fujisawa, D.M. Chudakov, P.M. Treuting, and A.Y. Rudensky. 2017. Stability and function of regulatory T cells expressing the transcription factor T-bet. Nature. 546:421–425. doi:10.1038/nature22360.

Lun, A.T.L., S. Riesenfeld, T. Andrews, T.P. Dao, T. Gomes, and J.C. Marioni. 2019. EmptyDrops: Distinguishing cells from empty droplets in droplet-based single-cell RNA sequencing data. Genome Biol. 20:1–9. doi:10.1186/S13059-019-1662-Y/FIGURES/3.

Mellanby, R.J., D. Thomas, J.M. Phillips, and A. Cooke. 2007. Diabetes in non-obese diabetic mice is not associated with quantitative changes in CD4+ CD25+ Foxp3+ regulatory T cells. Immunology. 121:15–28. doi:10.1111/J.1365-2567.2007.02546.X.

Muñoz-Rojas, A.R., and D. Mathis. 2021. Tissue regulatory T cells: regulatory chameleons. Nat. Rev. Immunol. 21:597–611. doi:10.1038/S41577-021-00519-W.

Nowakowska, D.J.D.J., and S. Kissler. 2016. Ptpn22 Modifies Regulatory T Cell Homeostasis via GITR Upregulation. J. Immunol. 196:2145–2152. doi:10.4049/jimmunol.1501877.

Ohnmacht, C., J. Park, S. Cording, J.B. Wing, K. Atarashi, Y. Obata, V. Gaboriau-Routhiau, R. Marques, S. Dulauroy, M. Fedoseeva, M. Busslinger, N. Cerf-Bensussan, I.G. Boneca, D. Voehringer, K. Hase, K. Honda, S. Sakaguchi, and G. Eberl. 2015. The microbiota regulates type 2 immunity through RORγt+ T cells. Science (80-.). 349:1–9. doi:10.1126/science.aac4263.

Pot, C., H. Jin, A. Awasthi, S.M. Liu, C.-Y. Lai, R. Madan, A.H. Sharpe, C.L. Karp, S.-C. Miaw, I.-C. Ho, and V.K. Kuchroo. 2009. Cutting Edge: IL-27 Induces the Transcription Factor c-Maf, Cytokine IL-21, and the Costimulatory Receptor ICOS that Coordinately Act Together to Promote Differentiation of IL-10-Producing Tr1 Cells. J. Immunol. 183. doi:10.4049/jimmunol.0901233.

Radichev, I.A., J. Yoon, D.W. Scott, K. Griffin, and A.Y. Savinov. 2020. Towards antigen-specific Tregs for type 1 diabetes: Construction and functional assessment of pancreatic endocrine marker, HPi2based chimeric antigen receptor. Cell. Immunol. 358. doi:10.1016/J.CELLIMM.2020.104224.

Robinson, M.D., and A. Oshlack. 2010. A scaling normalization method for differential expression analysis of RNA-seq data. Genome Biol. 11:R25. doi:10.1186/gb-2010-11-3-r25.

Sakai, R., M. Ito, K. Komai, M. Iizuka-Koga, K. Matsuo, T. Nakayama, O. Yoshie, K. Amano, H. Nishimasu, O. Nureki, M. Kubo, and A. Yoshimura. 2021. Kidney GATA3 + regulatory T cells play roles in the convalescence stage after antibody-mediated renal injury. Cell. Mol. Immunol. 18:1249– 1261. doi:10.1038/S41423-020-00547-X.

Schiering, C., T. Krausgruber, A. Chomka, A. Fröhlich, K. Adelmann, E.A. Wohlfert, J. Pott, T. Griseri, J. Bollrath, A.N. Hegazy, O.J. Harrison, B.M.J. Owens, M. Löhning, Y. Belkaid, P.G. Fallon, and F. Powrie. 2014. The alarmin IL-33 promotes regulatory T-cell function in the intestine. Nature. 513:564–568. doi:10.1038/nature13577.

Schuster, C., F. Jonas, F. Zhao, and S. Kissler. 2018. Peripherally induced regulatory T cells contribute to the control of autoimmune diabetes in the NOD mouse model. Eur. J. Immunol. doi:10.1002/eji.201847498.

Sefik, E., N. Geva-Zatorsky, S. Oh, L. Konnikova, D. Zemmour, A.M. McGuire, D. Burzyn, A. Ortiz-Lopez, M. Lobera, J. Yang, S. Ghosh, A. Earl, S.B. Snapper, R. Jupp, D. Kasper, D. Mathis, and C. Benoist. 2015. MUCOSAL IMMUNOLOGY. Individual intestinal symbionts induce a distinct population of RORgamma(+) regulatory T cells. Science (80-.). 349:993–997. doi:10.1126/science.aaa9420.

Simes, R.J. 1986. An improved Bonferroni procedure for multiple tests of significance. Biometrika. 73:751–754.

Tan, T.G., D. Mathis, and C. Benoist. 2016. Singular role for T-BET+CXCR3+ regulatory T cells in protection from autoimmune diabetes. Proc. Natl. Acad. Sci. U. S. A. 113:14103–14108. doi:10.1073/PNAS.1616710113.

Tenspolde, M., K. Zimmermann, L.C. Weber, M. Hapke, M. Lieber, J. Dywicki, A. Frenzel, M. Hust, M. Galla, L.E. Buitrago-Molina, M.P. Manns, E. Jaeckel, and M. Hardtke-Wolenski. 2019. Regulatory T cells engineered with a novel insulin-specific chimeric antigen receptor as a candidate immunotherapy for type 1 diabetes. J. Autoimmun. 103:102289. doi:10.1016/J.JAUT.2019.05.017.

Thornton, A.M., P.E. Korty, D.Q. Tran, E.A. Wohlfert, P.E. Murray, Y. Belkaid, and E.M. Shevach. 2010. Expression of Helios, an Ikaros transcription factor family member, differentiates thymic-derived from peripherally induced Foxp3+ T regulatory cells. J. Immunol. 184:3433–41. doi:10.4049/jimmunol.0904028.

Todd, J.A., M. Evangelou, A.J. Cutler, M.L. Pekalski, N.M. Walker, H.E. Stevens, L. Porter, D.J. Smyth, D.B. Rainbow, R.C. Ferreira, L. Esposito, K.M.D. Hunter, K. Loudon, K. Irons, J.H. Yang, C.J.M. Bell, H. Schuilenburg, J. Heywood, B. Challis, S. Neupane, P. Clarke, G. Coleman, S. Dawson, D. Goymer, K. Anselmiova, J. Kennet, J. Brown, S.L. Caddy, J. Lu, J. Greatorex, I. Goodfellow, C. Wallace, T.I. Tree, M. Evans, A.P. Mander, S. Bond, L.S. Wicker, and F. Waldron-Lynch. 2016. Regulatory T Cell Responses in Participants with Type 1 Diabetes after a Single Dose of Interleukin-2: A Non-Randomised, Open Label, Adaptive Dose-Finding Trial. PLOS Med. 13:e1002139. doi:10.1371/journal.pmed.1002139.

Watts, D., M. Janßen, M. Jaykar, F. Palmucci, M. Weigelt, C. Petzold, A. Hommel, T. Sparwasser, E. Bonifacio, and K. Kretschmer. 2021. Transient Depletion of Foxp3 + Regulatory T Cells Selectively Promotes Aggressive β Cell Autoimmunity in Genetically Susceptible DEREG Mice. Front. Immunol. 12. doi:10.3389/FIMMU.2021.720133.

Wicker, L.S., J. Clark, H.I. Fraser, V.E.S. Garner, A. Gonzalez-Munoz, B. Healy, S. Howlett, K. Hunter, D. Rainbow, R.L. Rosa, L.J. Smink, J.A. Todd, and L.B. Peterson. 2005. Type 1 diabetes genes and pathways shared by humans and NOD mice. J. Autoimmun. 25 Suppl:9–33. doi:10.1016/j.jaut.2005.09.009.

Wohlfert, E.A., J.R. Grainger, N. Bouladoux, J.E. Konkel, G. Oldenhove, C.H. Ribeiro, J.A. Hall, R. Yagi, S. Naik, R. Bhairavabhotla, W.E. Paul, R. Bosselut, G. Wei, K. Zhao, M. Oukka, J. Zhu, and Y. Belkaid. 2011. GATA3 controls Foxp3+ regulatory T cell fate during inflammation in mice. J. Clin. Invest. 121:4503–4515. doi:10.1172/JCI57456.

Xing, S., K. Gai, X. Li, P. Shao, Z. Zeng, X. Zhao, X. Zhao, X. Chen, W.J. Paradee, D.K. Meyerholz, W. Peng, and H.H. Xue. 2019. Tcf1 and Lef1 are required for the immunosuppressive function of regulatory T cells. J. Exp. Med. 216. doi:10.1084/jem.20182010.

Yamanouchi, J., D. Rainbow, P. Serra, S. Howlett, K. Hunter, V.E.S. Garner, A. Gonzalez-munoz, J. Clark, R. Veijola, R. Cubbon, S.-L. Chen, R. Rosa, A.M. Cumiskey, D. V Serreze, S. Gregory, J. Rogers, P.A. Lyons, B. Healy, L.J. Smink, J.A. Todd, L.B. Peterson, L.S. Wicker, P. Santamaria, E.S. Valerie, A. Gonzalez-munoz, J. Clark, R. Veijola, R. Cubbon, R. Rosa, A.M. Cumiskey, S. Gregory, J. Rogers, A. Paul, B. Healy, L.J. Smink, J.A. Todd, L.B. Peterson, S. Linda, and P. Santamaria. 2010. Interleukin-2 gene variation impairs regulatory T cell function and causes autoimmunity. Nat.Genet. 39:329–337. doi:10.1038/ng1958.Interleukin-2.

Yeh, W.I., H.R. Seay, B. Newby, A.L. Posgai, F.B. Moniz, A. Michels, C.E. Mathews, J.A. Bluestone, and T.M. Brusko. 2017. Avidity and Bystander Suppressive Capacity of Human Regulatory T Cells Expressing De Novo Autoreactive T-Cell Receptors in Type 1 Diabetes. Front. Immunol. 8. doi:10.3389/FIMMU.2017.01313.

Yu, F., S. Sharma, J. Edwards, L. Feigenbaum, and J. Zhu. 2015. Dynamic expression of transcription factors T-bet and GATA-3 by regulatory T cells maintains immunotolerance. Nat. Immunol. 16:197– 206. doi:10.1038/ni.3053.

Zhou, X., L.T. Jeker, B.T. Fife, S. Zhu, M.S. Anderson, M.T. McManus, and J.A. Bluestone. 2008. Selective miRNA disruption in T reg cells leads to uncontrolled autoimmunity. J. Exp. Med. 205:1983–91. doi:10.1084/jem.20080707.

Zhu, J., B. Min, J. Hu-Li, C.J. Watson, A. Grinberg, Q. Wang, N. Killeen, J.F. Urban, L. Guo, and W.E. Paul. 2004. Conditional deletion of Gata3 shows its essential function in TH1-TH2 responses. Nat. Immunol. 2004 511. 5:1157–1165. doi:10.1038/ni1128.

